# Hepatocyte-specific disruption of soluble epoxide hydrolase attenuates abdominal aortic aneurysm formation: novel role of the liver in aneurysm pathogenesis

**DOI:** 10.1101/2023.07.10.548127

**Authors:** David Kim, Tetsuo Horimatsu, Mourad Ogbi, Brandee Goo, Hong Shi, Praneet Veerapaneni, Ronnie Chouhaita, Mary Moses, Rosaria Prasad, Tyler W. Benson, Ragheb Harb, Ghaith Aboud, Hunter Seller, Steve Haigh, David J. Fulton, Gábor Csányi, Yuqing Huo, Xiaochun Long, Philip Coffey, Richard Lee, Avirup Guha, Darryl Zeldin, Sung Hee Hwang, Bruce D. Hammock, Neal L. Weintraub, Ha Won Kim

## Abstract

**Introduction:** Inflammation is a key pathogenic feature of abdominal aortic aneurysm (AAA). Soluble epoxide hydrolase (sEH) is a pro-inflammatory enzyme that converts cytochrome P450-derived epoxides of fatty acids to the corresponding diols, and pharmacological inhibition of sEH prevented AAA formation. Both cytochrome P450 enzymes and sEH are highly expressed in the liver. Here, we investigated the role of hepatic sEH in AAA using a selective pharmacological inhibitor of sEH and hepatocyte-specific Ephx2 (which encodes sEH gene) knockout (KO) mice in two models of AAA [angiotensin II (AngII) infusion and calcium chloride (CaCl_2_) application].

**Methods and results:** sEH expression and activity were strikingly higher in mouse liver compared with aorta and further increased the context of AAA, in conjunction with elevated expression of the transcription factor Sp1 and the epigenetic regulator Jarid1b, which have been reported to positively regulate sEH expression. Pharmacological sEH inhibition, or liver-specific sEH disruption, achieved by crossing sEH floxed mice with albumin-cre mice, prevented AAA formation in both models, concomitant with reduced expression of hepatic sEH as well as complement factor 3 (C3) and serum amyloid A (SAA), liver-derived factors linked to AAA formation. Moreover, sEH antagonism markedly reduced C3 and SAA protein accumulation in the aortic wall. Co-incubation of liver *ex vivo* with aneurysm-prone aorta resulted in induction of sEH in the liver, concomitant with upregulation of Sp1, Jarid1b, C3 and SAA gene expression, suggesting that the aneurysm-prone aorta secretes factors that activate sEH and downstream inflammatory signaling in the liver. Using an unbiased proteomic approach, we identified a number of dysregulated proteins [*e.g.,* plastin-2, galectin-3 (gal-3), cathepsin S] released by aneurysm-prone aorta as potential candidate mediators of hepatic sEH induction.

**Conclusion:** We provide the first direct evidence of the liver’s role in orchestrating AAA via the enzyme sEH. These findings not only provide novel insight into AAA pathogenesis, but they have potentially important implications with regard to developing effective medical therapies for AAA.

## Introduction

Aortic aneurysms are the 15th leading cause of death overall and the 10th in men older than age 55 in the United States, causing approximately 15,000 deaths per year in the US and more than 175,000 deaths globally (1, 2). Only ∼25% of patients with aortic rupture survive to surgery, so there is a pressing need to develop new approaches that can reduce abdominal aortic aneurysm (AAA) growth and rupture risk. Smoking, age, male sex, hypertension, and hypercholesterolemia are important risk factors for AAA, but the underlying mechanisms that lead to AAA formation are incompletely defined, which has hampered development of effective medical therapies.

Abdominal aortic aneurysms exhibit pronounced inflammation throughout the vascular wall, leading to tissue degeneration and progressive weakening of the aorta. Key pathways that orchestrate inflammation in AAA, however, are unknown. The liver amplifies systemic inflammation, and mounting evidence suggests that it may have an underappreciated role in the pathogenesis of AAA. For example, liver-derived molecules such as complement proteins (*e.g*., C3 and C5), fibrinogen, coagulation factors and serum amyloid A (SAA) have been linked to the pathogenesis of AAA (7-11). Moreover, clinical data suggest that AAA patients are at elevated risk for liver disease, while fatty liver disease is an independent risk factor for the development of AAA (12), further implying a potential linkage between the liver and AAA pathogenesis. Nevertheless, researchers have mainly focused on local vascular and inflammatory cells as mediators of aortic inflammation in AAA, while the potential role of the liver has largely been ignored.

Soluble epoxide hydrolase (sEH), encoded by the Ephx2 gene, is a widely distributed enzyme that catalyzes the hydrolysis of epoxy fatty acids (EpFA), such as epoxyeicosatrienoic acids [EETs, metabolized from arachidonic acid (AA)] produced by cytochrome P450 enzymes to their corresponding diols [i.e., dihydroxyeicosatrienoic acids (DHETs)]. These fatty acid-derived epoxides and diols are important lipid signaling molecules that regulate vascular inflammation. Pharmacological inhibition of sEH was shown to protect against hypertension and atherosclerosis (13-15), and limited data also suggested protection against AAA formation (15). However, neither the target cells nor the mechanisms of action of sEH inhibition were defined in AAA.

Expression studies in rodents and humans have determined that sEH, along with cytochrome P450 enzymes, are expressed highly in the liver as compared to other tissues (16, 17). Preferential expression of the sEH and the distantly regulated microsomal epoxide hydrolase (mEH) in the liver is consistent with the teleological role of epoxide hydrolase enzymes in xenobiotic metabolism (18). The very different substrate preference for these enzymes, however, suggests that the mEH is largely responsible for metabolism of reactive epoxides on cyclic backbones, with the sEH showing a high preference for EpFA. These findings raise the possibility that sEH expressed in liver could play a vital role in the pathogenesis of AAA. Here, using a pharmacological antagonist of sEH and liver-specific Ephx2 knockout (KO) mice, and two distinct models of AAA formation, we investigated the role of hepatic sEH in the pathogenesis of AAA. Our data show that either pharmacological sEH inhibition or liver-specific sEH gene disruption ameliorated the production of liver-derived inflammatory mediators linked to AAA and prevented AAA formation. In addition, we provide novel evidence that the aneurysm-prone aorta releases factors that can trigger induction of hepatic sEH and inflammatory mediators, suggesting a bidirectional circuit between the liver and aorta in the context of AAA.

## Materials and Methods

### Mice

Ephx2 floxed mice were obtained from NIEHS/NIH colony and bred with albumin-cre mice to create liver-specific Ephx2 KO (sEH^ΔLiver^) mice. For AngII infusion model, Ephx2^flox/flox^ mice and alb-Cre mice were bred with low density lipoprotein receptor knockout (LDLR KO) mice, respectively, and intercrossed to generate Ephx2^flox/flox^/alb-Cre mice in the LDLR KO background. Mice were anesthetized by isoflurane vaporizer (0.5-1.0 L/min for oxygen flowmeter, 4-5% for induction and 1-3% for maintenance, EZ Anesthesia Systems) and euthanized with intraperitoneal pentobarbital 150 mg/kg, inhaled anesthesia (isoflurane) followed by carbon dioxide (CO_2_) narcosis and cervical dislocation or bilateral thoracotomy, in accordance with AVMA Panel 2007 recommendations and institutional IACUC guidelines. All mice were randomized to different treatment groups to minimize experimental variability, and surgeons were blinded to group allocation to prevent accidental or selection bias. All animal experiments were performed at animal facilities of the Medical College of Georgia at Augusta University. Animal experimental protocols were approved by the Institutional Animal Care and Use Committee at the Medical College of Georgia at Augusta University and complied with National Institute of Health guidelines.

### AngII-induced AAA model

AngII (1,000 ng/kg/min, Enzo Life Sciences) was infused into 8 to 12 week old male mice via osmotic minipumps (ALZET Model 2004) as described previously (19, 20). In some experiments, mice were co-treated orally with an sEH inhibitor, TPPU [1-(1-Propionylpiperidin-4-yl)-3-(4-(trifluoromethoxy)phenyl)urea, 3-10 mg/kg body weight in drinking water] (21). Mice were euthanized at three weeks after minipump implantation, and abdominal aortic outer diameter was measured via microscopy. Aortic tissues, liver and blood were collected from mice for further analysis. AAA was defined as local dilation of the abdominal aorta that is at least 1.5 times the size of the reference aortic diameter.

### CaCl_2_-induced AAA model

CaCl_2_ application model of AAA (using 0.5 mol/L of CaCl_2_, Sigma-Aldrich) was performed as described previously (19, 20). In brief, following laparotomy, saline (sham control) or 0.5 mol/L of CaCl_2_ was applied to the infrarenal aortic adventitial surface for 15 min, followed by rinsing with 0.9% sterile saline and surgical closure. In some experiments, mice were co-treated orally with TPPU (3-10 mg/kg of body weight in drinking water) to inhibit sEH activity. After three weeks, mice were anesthetized, abdominal aortic outer diameter was measured via microscopy, and tissues were collected for further analysis.

### sEH activity assay

Soluble epoxide hydrolase activity was determined using sEH activity assay kit (BioVision) according to the manufacturer’s protocol. Briefly, liver and aorta tissues were homogenized with sEH assay buffer included in the kit and total protein was extracted. Protein concentration was determined with Pierce BCA Protein Assay Kit (ThermoFisher). sEH activity was calculated by measuring fluorescence (Ex/Em 362/460 nm) in kinetic mode using the standard curve and equation provided by manufacturer.

### Blood pressure measurement

Blood pressure was measured using a previously validated tail cuff method (Coda 6, Kent Scientific, Torrington, CT). Mice were conditioned to the instrument and procedure for 5 consecutive days prior to pump implantation. To insure a more robust estimation of systolic blood pressure (SBP), we used the interquartile mean of SBP measurements achieved through 30 measurement cycles every other day.

### Immunohistochemistry

Paraffin-embedded aortic tissue sections were stained with hematoxylin and eosin (H&E), Verhoeff van Gieson (VVG), Mac-3 (BD Pharmingen), MMP-2/-9 (Novus Biologicals), C3 (Abcam), SAA (R&D Systems), and HistoMouse-SP (Invitrogen) or DAB Substrate (Vector Labs) kits were used for visualization.

### Western Blotting

Protein extraction and Western blotting were performed as described previously (18, 19). Antibodies used were: sEH (Cayman Chemicals, 10010146), SAA (R&D Systems, AF2948), transferrin (Abcam, Ab82411), H3K4me3 (Cell Signaling Technology, 9751) and GAPDH (Invitrogen, AM4300).

### Quantitative PCR

Total RNA was extracted from aortic and liver tissues with QIAzol Lysis Reagent, and purified with RNeasy Lipid Tissue Mini Kit (Qiagen). Real time quantification of mRNA levels of the genes of interest was performed using Brilliant II SYBR Green QPCR Master Mix (Agilent Technologies) per manufacturer’s instructions. Normalized Ct values were subjected to statistical analysis and fold difference was calculated by ΔΔ Ct method as described previously (19, 20).

### *Ex vivo* co-incubation of liver with cigarette smoke extract

Fresh liver tissues harvested from 8 to 12 week old C57Bl/6 mice were cut into 3 to 5 mm slices (approximately 5 mg in weight) and co-incubated with 1% cigarette smoke extract (University of Kentucky Reference Cigarette 3R4F) in 500 µl of DMEM medium without FBS or antibiotics for 6 hr at 37°. Then, liver tissues were harvested for gene expression quantification as described above.

### *Ex vivo* co-incubation of liver with aorta

Following laparotomy in C57Bl/6 mice, saline (sham control) or 0.5 mol/L of CaCl_2_ was applied to the infrarenal aortic adventitial surface for 15 min, followed by rinsing with 0.9% sterile saline and surgical closure. After 7 days (a time point prior to AAA formation) (22), aortas were harvested. Liver tissues were freshly collected from control mice and cut into 3 to 5 mm slices, as described above, and co-incubated with two aortas harvested from sham- or CaCl_2_-treated mice in 500 µl of DMEM medium without FBS and antibiotics at 37°. After 12 hr of incubation, liver tissues were harvested to measure sEH activity and gene expression as described above.

### Liquid chromatography tandem mass spectrometry (LC-MS/MS) analysis, data processing and bioinformatics

Whole aortas from either sham- or CaCl_2_-treated mice were harvested 7 days after laparotomy and transferred to a 48-well plate with 500 µl of serum-free DMEM media. The conditioned media was collected after 12 hr incubation, centrifuged at 1000 g for 10 min, filtered through a Nalgene 0.2 µm pore vacuum filter, and concentrated using a 10 kDa MWCO Amicon Ultra-15 centrifugal filter (MiliporeSigma) operating at 1500 g for 30 min with 50 mM ammonium bicarbonate buffer until the final concentration was 2 mg/ml. Protein concentration was determined with Pierce BCA Protein Assay Kit (ThermoFisher). Protein digestion and mass spectrometry were performed as previously described (23). Briefly, extracted proteins were separated using an Ultimate 3000 nano-UPLC system (Thermo Scientific) and run on an Orbitrap Fusion Tribrid mass spectrometer (Thermo Scientific). Raw data were analyzed using Proteome Discoverer (v1.4, Thermo Scientific) and searched against the UniProt mouse protein database. Peptide-spectrum match (PSM) values, a total number of identified peptide spectra matched for the protein, were log-transformed to normalize the dataset. LIMMA package on R was used to compare protein levels in conditioned media of the aorta obtained from sham-versus CaCl_2_-treated mice, and p-values were adjusted using FDR (false discovery rate), with significance set at an adjusted p-value <0.05. The annotated genes were used as input for functional enrichment analysis, which was performed using the Gene Ontology (GO) terms.

### Statistical Analysis

Unless otherwise stated, all statistical analyses were performed using Graphpad Software (GraphPad Software, Inc., USA). Results are expressed as mean ± SEM. Differences between two groups were analyzed by student’s t-test. Multiple group datasets were evaluated for normality, and differences were analyzed by one-way ANOVA followed by Bonferroni post-hoc analysis. *p* values less than 0.05 were considered to be significant.

## Results

### sEH expression is significantly higher in the liver compared to aorta and further increased by AngII infusion

We first searched the available database (GeneAtlas U133A, gcrma) to check the expression levels of sEH in tissues and organs, and found that sEH is most highly expressed in liver compared to other tissues (**Supplemental Figure S1**). Our data likewise showed that sEH protein levels were > 10-fold higher in mouse liver compared to aorta as detected by Western blot (**Figure 1A**). Furthermore, sEH mRNA expression was significantly increased in livers, but not aortas, of AngII-infused mice (**Figure 1B**). sEH protein was also detected in the liver by immunofluorescence, which co-localized with the hepatocyte marker albumin and was increased by AngII infusion (**Figure 1C**). These findings are consistent with prior reports comparing cytochrome P450 enzyme and sEH expression in human and rodent tissues (16, 17).

**Figure 1.**
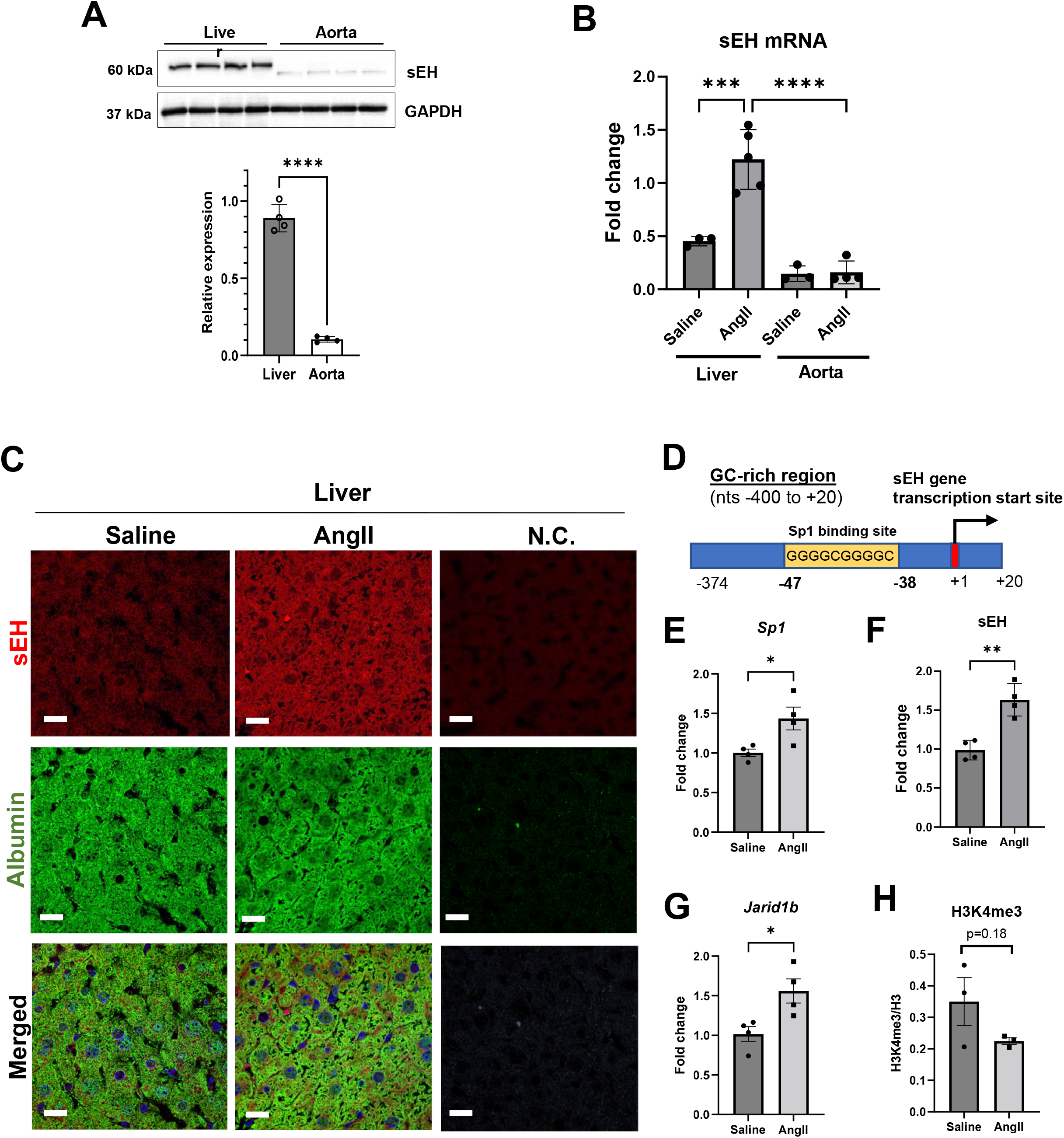
sEH expression is strikingly higher in the liver compared to aorta and further increased by AngII infusion. (**A**) Comparison of sEH protein expression between liver and aorta as determined by Western blot. (**B**) sEH mRNA expression in liver of saline controls and AngII-infused mice. (**C**) Immunofluorescence analysis of sEH in liver tissues of AngII-infused mice. Albumin was used as a marker for hepatocytes. Scale bar = 20 µm. (**D**) Depiction of the Sp1 promoter binding site in GC-rich region between nucleotides -374 and +28 with respect to the putative sEH gene transcription start site. mRNA expression of Sp1 (**E**), sEH (**F**), and Jarid1b as determined by qPCR (**G**) and quantification of H3K4me3 levels detected by Western blot (**H**). Data are represented as mean ± SEM. *p<0.05, **p<0.01, ****p<0.0001.

We subsequently investigated the molecular mechanisms of hepatic sEH gene induction during AngII-induced AAA. Sp1 was previously reported as a major transcription factor for sEH gene expression in various cell lines, including human liver cancer-derived HepG2 cells (24). The minimal promoter was identified as a GC-rich region between nucleotides -374 and +28 with respect to the putative transcription start site (**Figure 1D**). Sp1 expression was markedly increased in livers of AngII-infused mice compared to saline control (**Figure 1E**), in conjunction with elevated sEH expression (**Figure 1F**). Histone H3K4 demethylase Jarid1b was also reported to positively regulate sEH expression in response to AngII (25), and our data confirmed upregulated Jarid1b expression (**Figure 1G**), along with a trend toward reduced H3K4me3 (indicator of Jarid1b activity) (**Figure 1H**), in livers of mice infused with AngII. These findings suggest that Sp1 and/or Jarid1b may mediate induction of hepatic sEH in the context of AngII-induced AAA formation.

Smoking, male sex and aging are major risk factors for AAA, and hepatic sEH expression was reported to be positively correlated with age and male sex (26, 27). Moreover, smoke exposure has been shown to stimulate lung sEH activity and inflammation in mice (28). To investigate whether smoke exposure may be associated with induction of sEH and inflammatory gene expression in liver tissue, we treated mouse liver slices with 1% cigarette smoke extract (CSE) for 6 hr. Interestingly, CSE induced hepatic sEH, and inflammatory mediators SAA and C3 (both of which have been linked to AAA) (7-11), in mouse liver (**Supplemental Figure S2**).

### Pharmacological inhibition of sEH reduced sEH activity in the liver, but not in aorta, and protected against AngII-induced AAA formation

Next, we quantified sEH activity in the liver and aorta in the murine AngII-induced AAA model. Infusion of AngII into LDLR KO mice to induce AAA resulted in increased sEH activity in the liver, which was reduced by concurrent treatment with a highly selective and potent sEH inhibitor, TPPU, (3-10 mg/kg of BWT in drinking water) (21). In contrast, aortic sEH activity was virtually undetectable, in the presence or absence of TPPU (**Figure 1A**). To investigate whether sEH inhibition could mitigate AAA formation, we administered TPPU orally to LDLR KO mice, followed by AngII infusion. TPPU treatment (3 mg or 10 mg/kg body weight) significantly reduced AAA incidence (**Figure 2B**) and aortic diameter (**Figure 2C**) without affecting systolic blood pressure (**Supplemental Figure S3**). Representative *in situ* abdominal aortic images of each group are shown in **Figure 2D**. Prominent elastin fragmentation (H&E and VVG staining) and macrophage infiltration (Mac-3 staining) were observed in aortas of AngII-infused mice, which were markedly inhibited by TPPU treatment (**Figure 2E**). Furthermore, TPPU markedly decreased expression of matrix metalloproteinase (MMP)-2/-9 in the aorta (**Figure 2E**).

**Figure 2.**
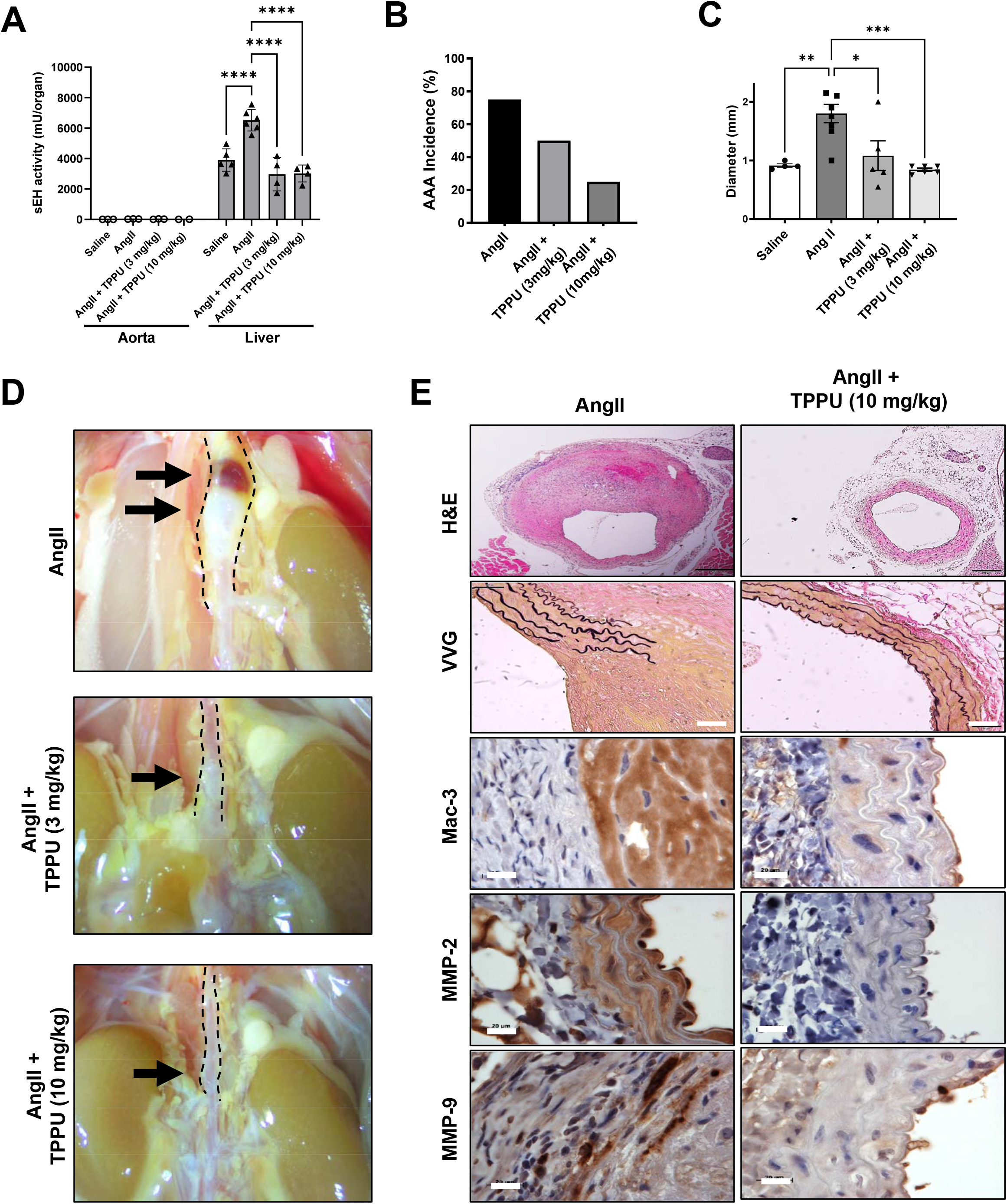
Pharmacological inhibition of sEH reduced sEH activity in the liver, but not in aorta, and protected against AngII-induced AAA formation. AngII was infused for 3 weeks via osmotic minipump in LDLR KO mice with or without TPPU treatment. (**A**) sEH enzymatic activity in liver and aorta. (**B**) AAA incidence. (**C**) Aortic diameter. (**D**) Representative *in situ* images of aorta in AngII-infused mice with or without TPPU treatment; dashed lines delineate aorta. (**E**) Representative images of histology, H&E, VVG (elastin degradation, scale bar = 50 µm), Mac-3 (macrophages) and MMP-2/-9 (scale bar = 20 µm). Data are represented as mean ± SEM. *p<0.05, **p<0.01, ***p<0.001, ****p<0.0001.

### sEH inhibitor blunted expression of inflammatory mediators, ER stress genes and macrophage infiltration in the liver of AngII-infused mice

To investigate the impact of sEH inhibition on inflammatory mediator production in the liver, we quantified the expression of complement factors and SAA, previously reported to contribute to AAA formation (7, 8, 11), in the livers of mice infused with AngII. Interestingly, expression of C3 and C3aR was significantly increased by AngII infusion and reduced by TPPU treatment (**Figure 3A&B**). IL-6 expression, reportedly associated with liver inflammation (29), was also markedly increased in the livers of AngII-infused mice and reduced by TPPU treatment (**Figure 3C**). Furthermore, TPPU reduced SAA mRNA expression in liver (**Figure 3D**), and SAA protein content in liver, plasma and aorta (**Figure 3E&F**). Given that SAA mRNA expression was very low in aorta compared to liver (**Supplemental Figure S4**), SAA protein that accumulated in the aorta was likely derived from circulating SAA secreted from the liver. There was no change in hepatic expression of fibrinogen α, β or γ, which have also been reported to be associated with AAA formation (9), in response to AngII infusion or TPPU treatment (**Supplemental Figure S5**).

**Figure 3.**
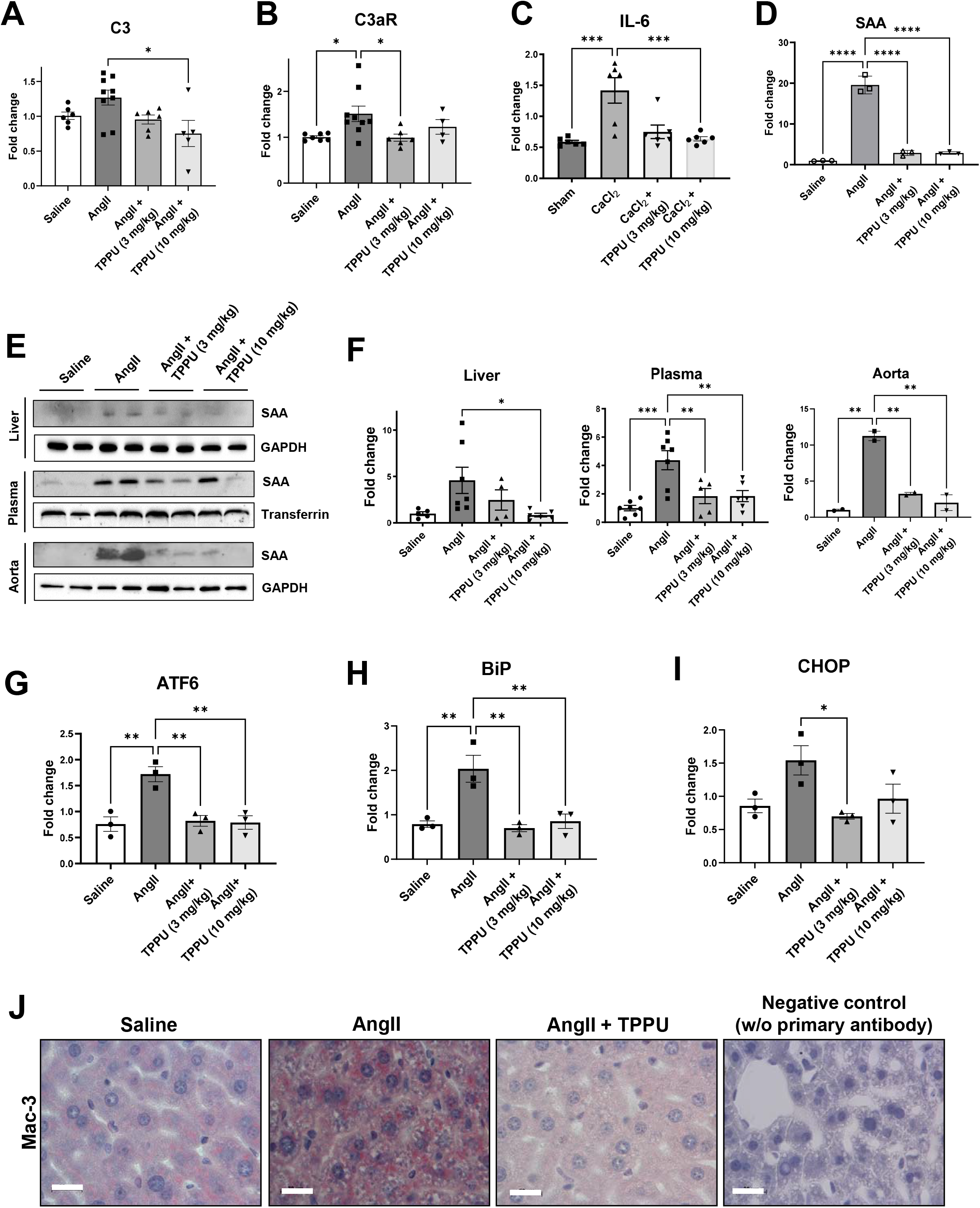
sEH inhibitor blunted expression of inflammatory mediators, ER stress genes and macrophage infiltration in the liver of AngII-infused mice. AngII was infused for 3 weeks via osmotic minipump in LDLR KO mice with or without TPPU treatment. mRNA expression of C3 (**A**), C3aR (**B**), IL-6 (**C**) and SAA (**D**) in liver tissues. SAA protein expression in liver, plasma and aortic tissues (**E**) and densitometric analysis (**F**). mRNA expression of ATF (**G**), BiP (**H**) and CHOP (**I**) in liver tissues. (**J**) Immunohistochemical staining of Mac-3 (macrophages) in liver tissues. Scale bar = 20 µm. Data are represented as mean ± SEM. *p<0.05, **p<0.01, ***p<0.001, ****p<0.0001.

Endoplasmic reticulum (ER) stress plays an important role in liver inflammation and stress signaling (30). Expression of hepatic ER stress gene markers, ATF6, BiP and CHOP, was significantly increased in the livers of AngII-infused mice and blunted by TPPU treatment (**Figure 3G-I**). Furthermore, AngII infusion increased macrophage infiltration in the liver, which was attenuated by TPPU treatment (**Figure 3J**). Taken together, these results support the hypothesis that liver inflammatory responses temporally correlate with AAA formation in mice and are mitigated by sEH inhibition.

### Hepatocyte-specific sEH gene disruption protected against AngII-induced AAA formation in conjunction with reduced expression of liver-derived inflammatory mediators

Although TPPU is a highly selective and potent inhibitor of sEH, we cannot exclude the possibility of off-target effects, and it does not selectively target sEH in the liver. Thus, we created liver-specific sEH deficient mice by crossing Ephx2 floxed mice (Ephx2^flox/flox^) and albumin (alb)-Cre mice. The presence of WT/flox Ephx2 allele and/or alb-Cre allele in mice was validated via PCR-based genotyping (**Supplemental Figure S6A**). We confirmed that Ephx2 gene expression is diminished in liver, but not aorta, in Ephx2^flox/flox^/alb-Cre mice (**Supplemental Figure S6B**). Notably, liver-specific sEH gene disruption significantly reduced AAA incidence (**Figure 4A**) and aortic diameter (**Figure 4B**) in AngII-infused mice, without affecting blood pressure (**Supplemental Figure S7**), in conjunction with reduced SAA and C3 expression in liver (**Figure 4C&D**). Liver-specific Ephx2 KO mice also exhibited reduced elastin degradation, macrophage infiltration, and MMP2/MMP9 expression (**Figure 4E**). Furthermore, SAA and C3 protein accumulation in the aortic wall was ameliorated by liver-specific sEH gene disruption (**Figure 4F**).

**Figure 4.**
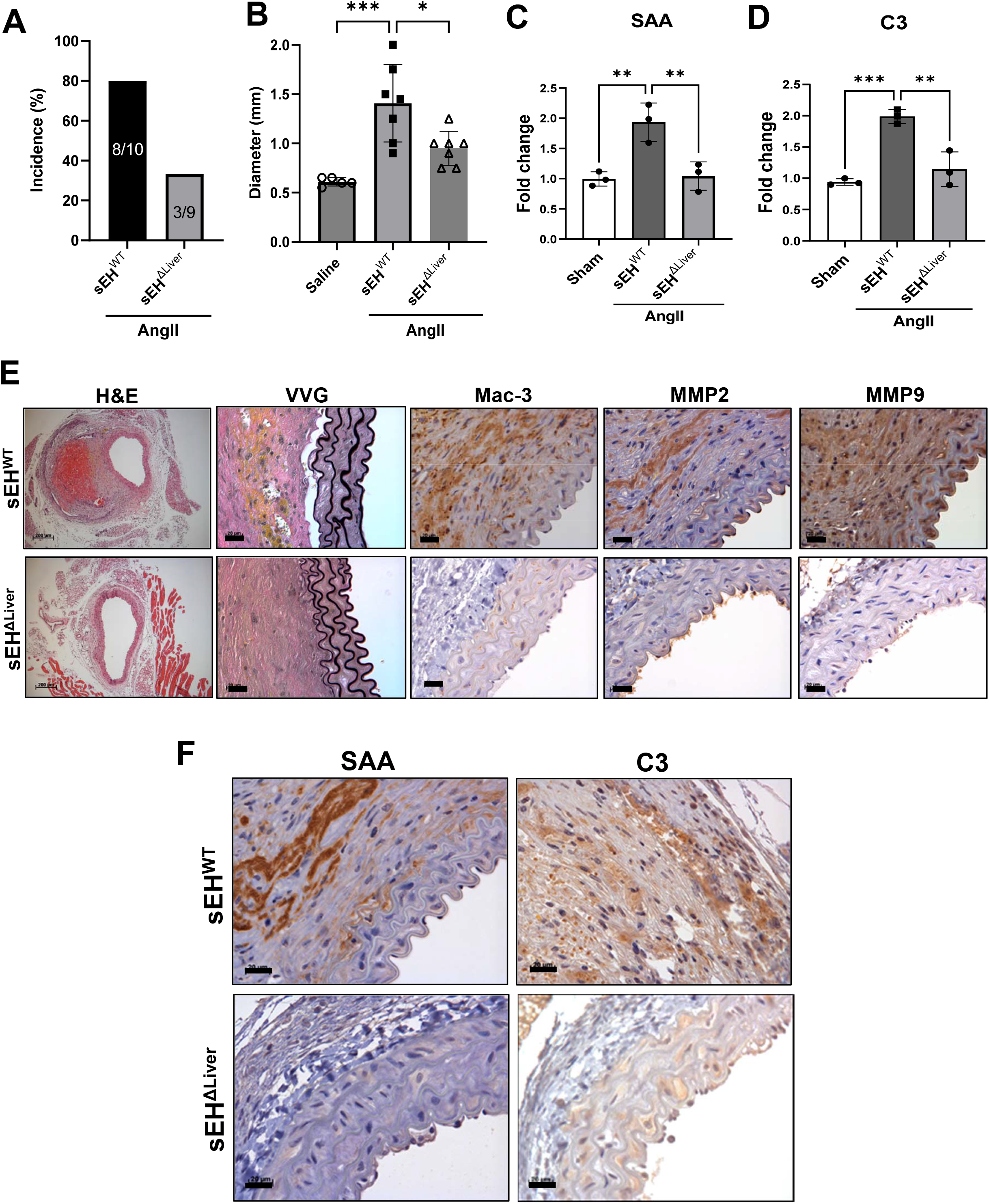
Hepatocyte-specific sEH gene disruption protected against AngII-induced AAA formation in conjunction with reduced expression of liver-derived inflammatory mediators. AngII was infused via osmotic minipump in littermate WT (sEH^WT^) and hepatocyte-specific sEH deficient mice (sEH^ΔLiver^) in the LDLR KO background. (**A**) AAA incidence. (**B**) Aortic diameter. (**C**) SAA mRNA expression. (**D**) C3 mRNA expression. (**E**) Representative histology images of H&E, VVG (elastin degradation), Mac-3 (macrophages) and MMP-2/-9. (**F**) Immunohistochemical analysis for SAA and C3 protein accumulation in aortic wall of AngII-infused mice. Scale bar = 20 µm. Data are represented as mean ± SEM. *p<0.05, **p<0.01, ***p<0.001.

### CaCl_2_-induced AAA model: sEH expression and effects of TPPU

Next, we investigated whether hepatic sEH expression is also induced by aortic CaCl_2_ application, which is a local acute aortic injury model with a well-defined time course (22). Consistent with data in the AngII infusion model, sEH protein expression was markedly increased in the livers of CaCl_2_-treated mice compared to control (**Figure 5A**). In addition, there was a trend toward increased expression of SAA in liver, plasma and aorta, and reduction by TPPU treatment, following aortic CaCl_2_ application (**Supplemental Figure S8**). Notably, TPPU treatment likewise protected against AAA formation induced by CaCl_2_ application (**Figure 5B&C**), in conjunction with decreased elastin degradation, macrophage infiltration and MMP-2/-9 expression (**Figure 5D**).

**Figure 5.**
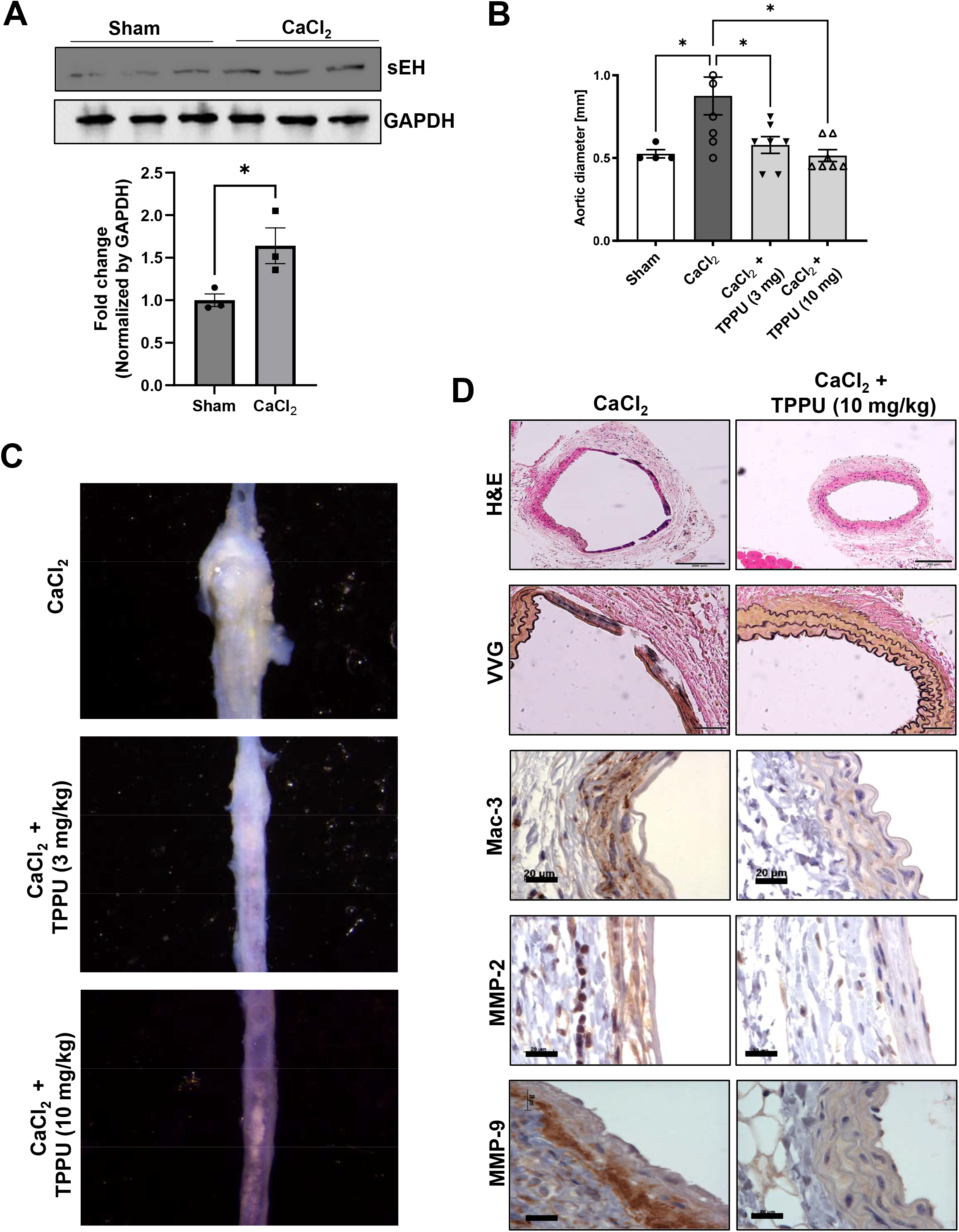
sEH expression is increased in the livers of CaCl_2_-induced AAA mice and pharmacological inhibition of sEH protects against AAA formation. CaCl_2_ or saline (sham) was applied to the infrarenal aorta of C57Bl/6 mice with or without TPPU treatment. (**A**) sEH protein expression in liver as measured by Western blot. (**B**) Aortic diameter. (**C**) Representative images of aorta in mice with or without oral TPPU treatment. (**D**) Representative histology; H&E, VVG (elastin degradation), Mac-3 (macrophages) and MMP-2/-9. Scale bar = 20 µm. Data are represented as mean ± SEM. *p<0.05.

### Liver-specific sEH gene disruption protected against CaCl_2_-induced AAA formation in conjunction with reduced expression of liver inflammatory mediators

Next, we employed liver-specific Ephx2 KO mice in the CaCl_2_ application model. Liver-specific disruption of sEH strongly protected against CaCl_2_-induced AAA formation (**Figure 6A&D**), in association with reduced hepatic expression of SAA and C3 (**Figure 6B&C**). Liver-specific s Ephx2 gene disruption also ameliorated major pathological features of AAA, such as elastin degradation (VVG), macrophage infiltration (Mac-3) and MMP-2/-9 expression (**Figure 6E**). SAA and C3 protein accumulation in the aorta was also significantly reduced in liver-specific sEH KO mice compared to WT mice (**Figure 6F**).

**Figure 6.**
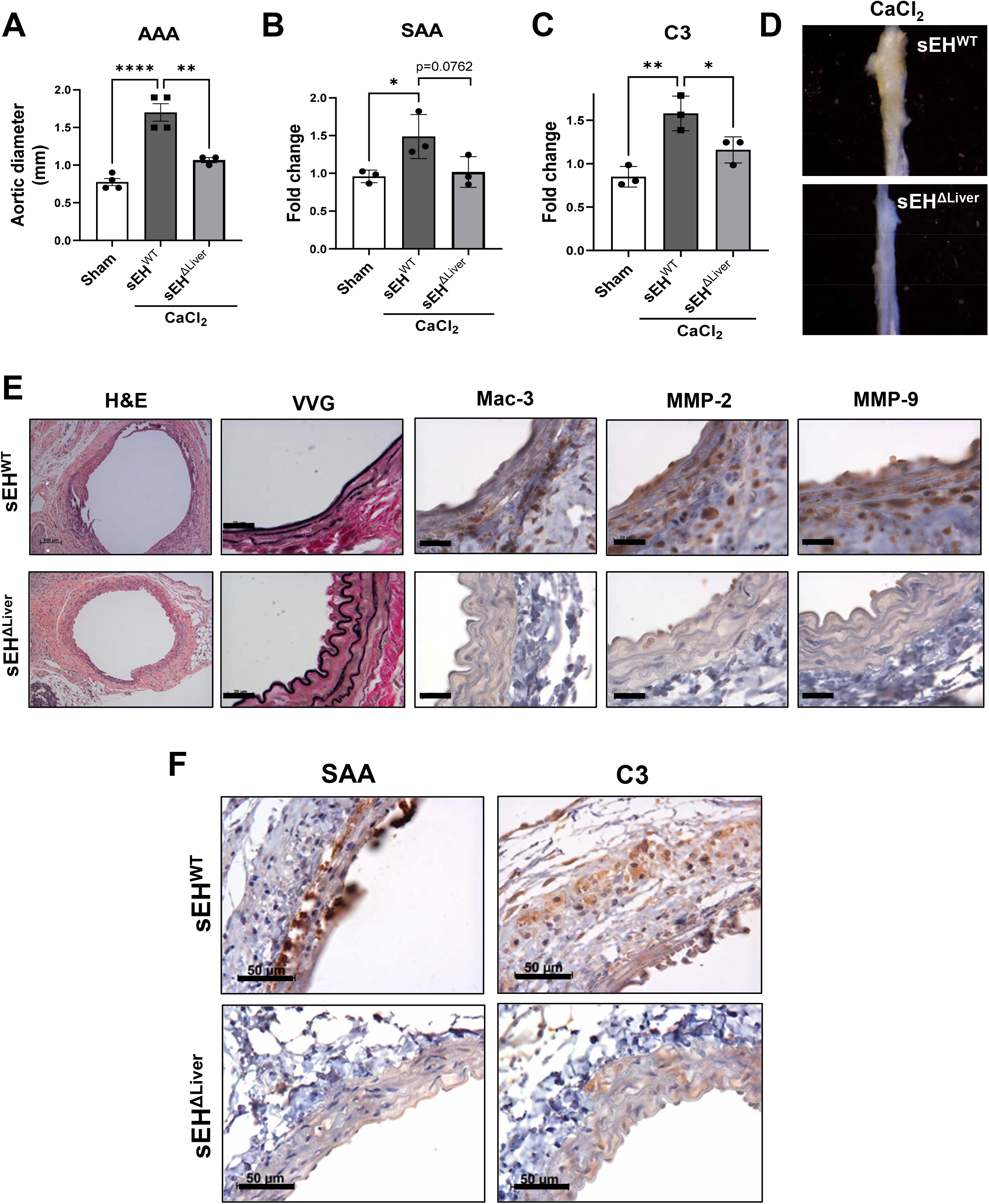
Liver-specific sEH gene disruption protected against CaCl_2_-induced AAA formation in conjunction with reduced expression of liver inflammatory mediators. CaCl_2_ or saline (sham) was applied to the infrarenal aorta of littermate WT (sEH^WT^) and hepatocyte-specific sEH deficient mice (sEH^ΔLiver^). (**A**) Aortic diameter. (**B**) SAA mRNA expression. (**C**) C3 mRNA expression. (**D**) Representative aortic images of aorta in sEH^WT^ and sEH^ΔLiver^ mice. (**E**) Representative images of H&E, VVG (elastin degradation), Mac-3 (macrophages) and MMP-2/-9. Scale bar = 20 µm. (**F**) Immunohistochemical analysis of SAA and C3 protein accumulation in the aortic wall. Scale bar = 50 µm. Data are represented as mean ± SEM. *p<0.05, **p<0.01, ****p<0.0001.

### Co-incubation of CaCl_2_-treated aorta with liver augmented hepatic sEH gene transcription and inflammatory gene expression

Our data in the CaCl_2_ model suggested that local aortic injury might trigger secretion of molecules that can induce hepatic sEH expression and inflammatory mediator production. To test this hypothesis, we co-incubated sham- or CaCl_2_-treated, aneurysm-prone aorta (harvested 7 days after laparotomy, prior to AAA formation) with liver harvested from control mice for 12 hr (**Figure 7A**). Compared to sham aorta, CaCl_2_-treated aorta significantly increased sEH gene expression and activity in the liver (**Figure 7B&C**), in conjunction with increased C3 & SAA expression (**Figure 7D&E**). We identified Sp1 and Jarid1b as potential upstream regulators of sEH induction in AngII-infused mice (**Figure 2F-J**) (24, 25). Likewise, we observed that compared with sham aorta, co-incubation with CaCl_2_-treated aorta significantly upregulated the expression of Sp1 and Jarid1b in liver tissue (**Figure 7F&G**).

**Figure 7.**
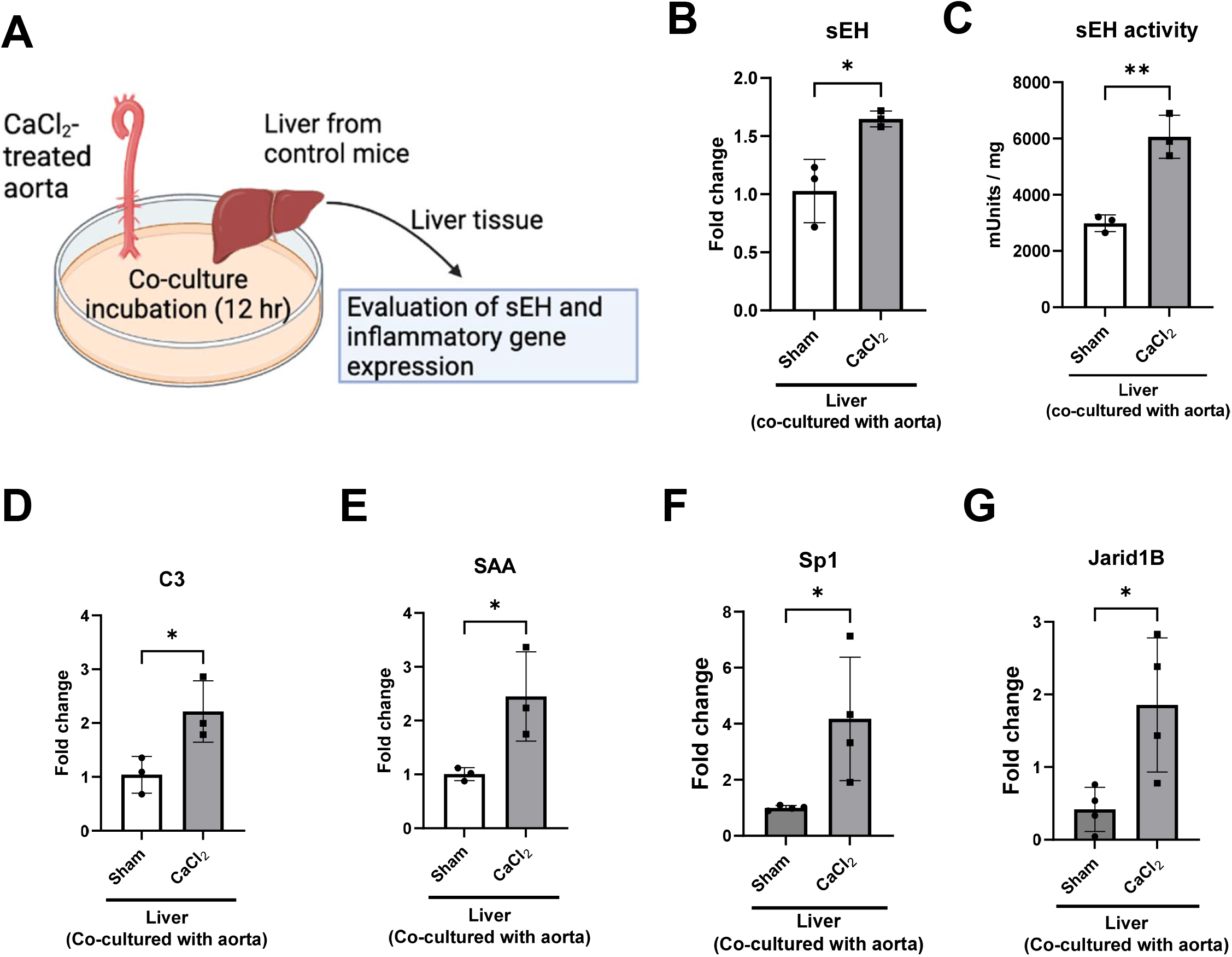
Co-incubation of CaCl_2_-treated aorta with liver augmented hepatic sEH gene transcription and inflammatory gene expression. Fresh liver slices from control mice were co-incubated for 12 hr with aorta harvested from sham- or CaCl_2_-treated mice 7 days after laparotomy. (**A**) Experimental scheme. sEH mRNA expression (**B**) and enzymatic activity (**C**) in liver tissues. Hepatic mRNA expression of Sp1 (**D**), Jarid1b (**E**), C3 (**F**) and SAA (**G**). Data are represented as mean ± SEM. *p<0.05, **p<0.01.

### Profiling of candidate molecules secreted from the aneurysm-prone aorta

To begin to identify signaling mediators released from aneurysm-prone aorta that might induce hepatic sEH, we performed LC-MS/MS-based unbiased proteomic analysis of incubation medium collected from sham- or CaCl_2_-treated aorta (**Figure 8A**). We identified a number of upregulated proteins [*e.g.,* plastin-2, kininogen-1, prosaposin, periostin, calreticulin, galectin-3 (gal-3), cathepsin S and xanthine dehydrogenase/oxidase] and downregulated proteins (*e.g.,* fatty acid synthase, glycerol-3-phosphate dehydrogenase, transketolase, troponin C, glutamine synthase and citrate synthase) in incubation media from CaCl_2_-treated aorta compared to sham (**Figure 8B-D**). Interestingly, GO enriched analysis predicted the upregulation in cell spreading, exocytosis and apoptotic signaling pathways (**Supplemental Figure S9**), and downregulation in NAD^+^/nicotinamide metabolism and glycolytic process pathways (**Supplemental Figure S10**) in CaCl_2_-treated compared to control aorta.

**Figure 8.**
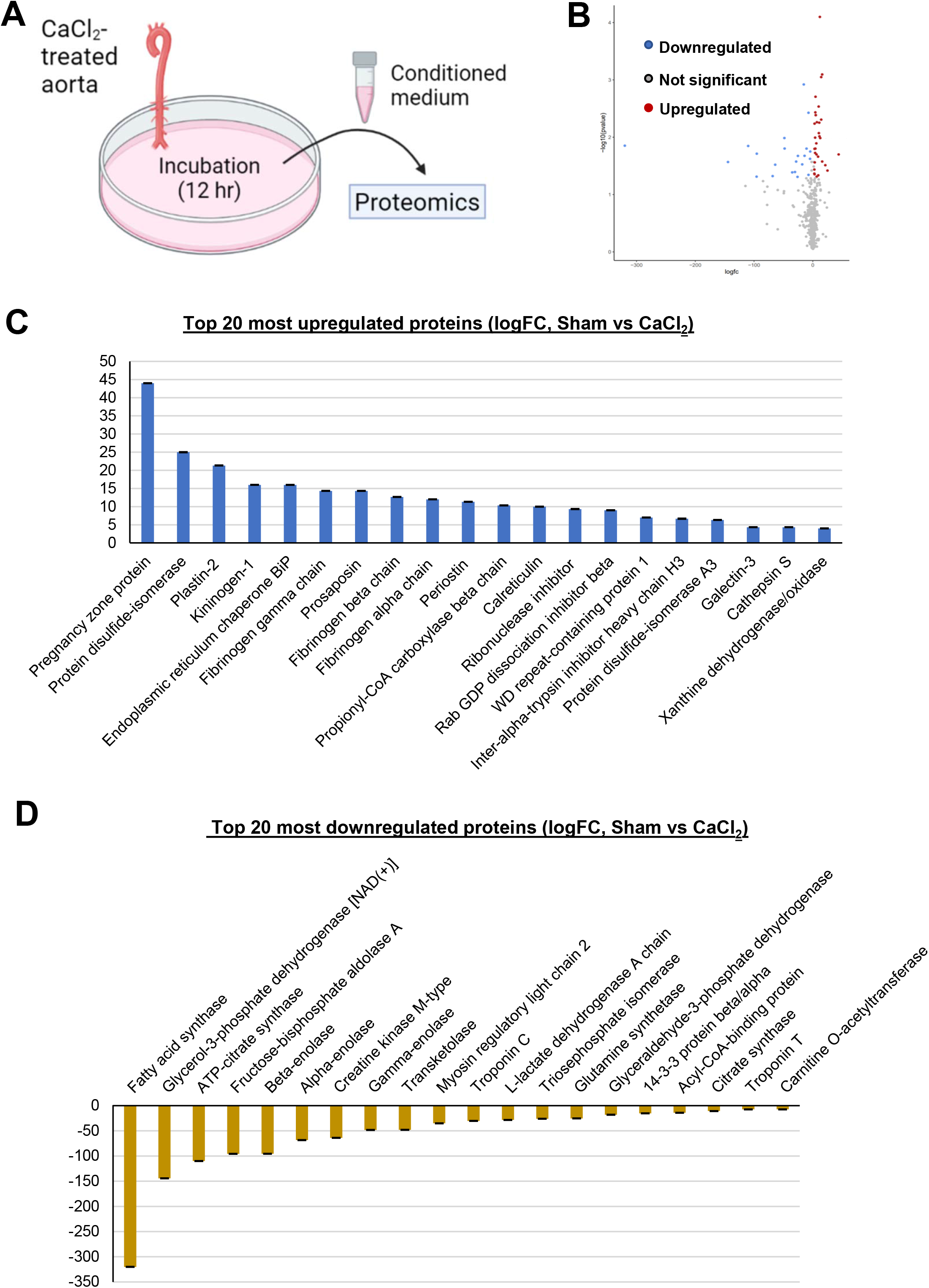
Profiling of candidate molecules secreted from the aneurysm-prone aorta. Whole aortas from sham- or CaCl_2_-treated mice harvested 7 days after laparotomy were incubated for 12 hr, and conditioned media were then collected for proteomic analysis. (**A**) Experimental scheme. (**B**) Volcano plot of all differentially-expressed proteins between sham and CaCl_2_ treatment. Top 20 most upregulated (**C**) and downregulated (**D**) proteins as predicted by gene ontology (GO) enriched analysis.

## Discussion

The liver is widely accepted to play a crucial role in atherosclerosis, which like AAA is a vascular inflammatory disease (3). Moreover, liver-targeted therapeutics (*e.g*., statins and PCSK9 inhibitors) are highly effective at preventing and treating atherosclerosis (4, 5). Here, using a pharmacological inhibitor and liver-specific gene disruption approach in two distinct animal models of AAA, we tested the hypothesis that sEH expressed specifically in the liver plays a crucial role in AAA formation. Major findings of this study are: 1) sEH expression and activity in the liver, but not aorta, is upregulated in the context of AAA formation, potentially through induction of the transcription factor Sp1 and/or the histone demethylase Jarid1b; 2) treatment with a pharmacological sEH inhibitor prevented AAA formation in conjunction with diminished expression of liver-derived inflammatory mediators, 3) accumulation of liver-derived pro-inflammatory mediators in the aorta is prevented by pharmacological sEH inhibition or liver-specific sEH gene disruption, 4) liver-specific Ephx2 KO mice are protected against AAA formation; 5) aneurysm-prone aorta releases transferrable factors that trigger induction of sEH in the liver. These findings provide novel insight into the liver’s role in orchestrating AAA via sEH and may have important implications with regard to developing effective medical therapy for this disorder.

sEH is widely expressed in various cell types and tissues including liver, kidney, muscle and vascular cells such as vascular smooth muscle cells (16), and pharmacological inhibition of sEH has been shown to improve vascular function and diminish inflammation. Limited data also suggest that pharmacological inhibition of sEH protects against AAA formation (15). Effects of sEH inhibitors on the vasculature are generally presumed to be due to local inhibition of vascular EET metabolism, as EETs that have powerful vasodilatory and anti-inflammatory effects are produced in blood vessels. However, sEH expression is by far highest in the liver compared to other tissues and organs (16, 17), and hepatic sEH expression is upregulated with aging, smoking and male sex (24-26), all of which are important risk factors for AAA. The liver also has high specific activities of the cytochrome P450 enzymes which, among other functions, generate the inflammation-resolving epoxy fatty acids. The liver is a very large tissue and heavily vascularized by the sinusoid system. Thus, it should dominate both the production of epoxy fatty acids and their hydrolysis to diols at the whole-body level. A dominant role of the liver in producing EpFA and their hydrolysis products was recently reported in the context of therapy for traumatic brain injury (31) and Alzheimer’s disease (32). In general, the EpFA appear to be inflammation-resolving lipid mediators while the diols, especially the linoleate diols, lack this activity and may even be proinflammatory (33).

The liver can amplify systemic inflammation via a number of biochemical pathways. Complement proteins, synthesized by the liver and circulating in the blood, are well-defined acute phase proteins associated with systemic inflammation. Complement proteins participate in recruitment of immune cells and activation of coagulation pathways, both of which have a potential role in the initiation and progression of AAA (7, 8). Importantly, complement depletion protected against AAA via reducing neutrophil recruitment, while activation of C3 or C5 promoted AAA formation (7). Furthermore, the complement regulator CD59 protected against AngII-induced AAA formation (34). In addition, complement factors and their degradation products were reported to be highly expressed in human AAA (10), further suggesting an important role for liver-derived complement factors in AAA pathogenesis. SAA is another major inflammatory factor produced primarily in the liver, and its levels in blood are increased in various chronic inflammatory diseases including obesity (35) and diabetes (36). SAA was also reported to be elevated by smoking (37), a major risk factor for AAA, and SAA-deficient mice were reported to be resistant to AAA formation (11), suggesting that SAA derived from liver could function as a key mediator in the pathogenesis of AAA. In both the AngII-infusion and CaCl_2_ models of AAA formation, C3 and SAA gene expression was significantly upregulated in the liver and abrogated by liver-specific sEH disruption. Furthermore, C3 and SAA protein accumulation in the aortic wall was markedly attenuated by liver-specific sEH gene disruption, suggesting that sEH in the liver controls the production of key inflammatory mediators that circulate from the liver and contribute to the pathogenesis of AAA. However, the complete repertoire of mediators regulated by sEH in the liver, and how these mediators (individually or collectively) promote AAA, remains to be defined.

Hepatic sEH expression was upregulated in both the AngII infusion and CaCl_2_ models of AAA formation. The sEH enzyme is often considered both a cause of inflammation by degrading inflammation-resolving EpFA and a marker of inflammation due to its increased transcription and translation (38). AngII was reported to induce sEH via the histone H3K4 demethylase Jarid1b in murine aorta, (25). The mechanism, however, was determined to be independent of Jarid1b’s histone demethylase activity. Rather, Jarid1b was detected to bind to the 3’UTR region of Ephx2 mRNA, thereby maintaining UTR length and increasing its stability (25). Additionally, Sp1 was reported as a major transcription factor for sEH gene expression (24). Both Sp1 and Jarid1b expression were upregulated in livers of mice with AngII-induced AAA compared to saline control, suggesting a potential mechanistic link between AAA induction and hepatic sEH expression. Further studies will be required to determine the specific role of Sp1 and/or Jarid1b in the upregulation of sEH in the liver and isolated hepatocytes, and the potential relevance to AAA formation.

The CaCl_2_ model of AAA formation is considered to be a local acute aortic injury model of AAA. We co-incubated aortas collected from mice 7 days after CaCl_2_ application (a time point that precedes formation of AAA) with mouse liver *ex vivo* and provide evidence that the aneurysm-prone aorta is capable of secreting factors that induce hepatic sEH. Moreover, this induction of sEH is associated with upregulation of Sp1 and Jarid1b expression, similar to that observed in livers of mice during experimental AAA formation *in vivo*. Using an unbiased proteomics approach, we identified molecules that were differentially expressed (up- or down-regulated) in the conditioned media collected from aneurysm-prone aorta compared to sham control. Damage-associated molecular patterns (DAMPs) [*e.g.*, high-mobility group box 1 (HMGB1), osteopontin (Spp1), galectin-3 (gal-3)] are endogenous signaling molecules released from damaged or dying cells that activate the immune system. Intriguingly, HMGB1, Spp1 and gal-3 were reported to be causally linked to both AAA (39-42) and liver disease (43-45). Among those DAMPS, only gal-3 was significantly upregulated in conditioned media from CaCl_2_-treated aorta, while there was no change in HMGB1 or Spp1. Plastin is a critical regulator of actin dynamics in cells of both the adaptive and innate immune systems. Interestingly, plastin-2 expression was increased more than 20-fold in the CaCl_2_-treated group compared to sham control in our proteomics data. Moreover, secretion of cathepsin S, a pro-inflammatory protease, and xanthine dehydrogenase/oxidase, a reactive oxygen species-generating enzyme, was also significantly increased by ∼4 fold in the CaCl_2_-treated group. In contrast, glycerol-3-phosphate dehydrogenase, a protein related to nicotinamide adenine dinucleotide (NAD+) metabolism, was found to be significantly downregulated by CaCl_2_ treatment. Collectively, these data demonstrate dysregulated secretion of many proteins by aneurysm-prone aorta, some of which are plausibly linked to AAA formation. Further investigations will be required, however, to identify those factors responsible for induction of hepatic sEH expression and inflammatory mediators in the context of AAA.

sEH inhibitors have proven safe and been tested in several clinical trials, including for chronic pain. Three sEH inhibitors, AR9281 (46), GSK2256294 (47), and EC5026 (48) have successfully completed Phase Ia trials with no significant safety concerns. TPPU, employed in this study, was a hit to develop EC5026 (49) and the most studied sEH inhibitor in the preclinical studies; similar in structure, equally potent in mice and rats to sEH inhibitors under study in humans (50), and TPPU metabolism in rats and humans is very similar, suggesting that data obtained from mice in this study could potentially be extrapolated to humans. Our findings suggest that sEH inhibitors currently being evaluated in several clinical trials potentially could be repurposed to treat AAA.

A limitation of this study is that commercially available assay kits to detect sEH activity could potentially also detect other enzymes with hydrolase activities, such as glutathione S transferases, which may also open epoxide groups and release the fluorophore detected by these kits. More specific sEH activity assay kits are currently under development, which should lead to more accurate detection of sEH activity in the future.

In conclusion, we report that sEH present in the liver plays a crucial role in the pathogenesis of AAA. We also provide evidence of a novel bidirectional circuit between aorta and liver in the context of AAA, wherein the aneurysm-prone aorta secretes factors that trigger the liver to produce inflammatory mediators, which in turn contribute to AAA. Pharmacological inhibitors of sEH, or gene therapy targeting sEH in the liver, may represent attractive strategies to prevent or treat AAA.

## Abbreviations and Acronyms

AA: arachidonic acid
AAA: abdominal aortic aneurysms
Abl: albumin
AngII: angiotensin II
CaCl2: calcium chloride
CVD: cardiovascular disease
DAMP: damage-associated molecular pattern
DHETs: dihydroxyeicosatrienoic acids
EETs: epoxyeicosatrienoic acids
ER: Endoplasmic reticulum
Gal-3: galectin-3
HMGB1: high-mobility group box 1
KO: knockout
MMP: matrix metallopeptidase
SAA: serum amyloid A
sEH: soluble epoxide hydrolase
Spp1: osteopontin
VVG: van Gieson stain

