## Supplemental for "Hepatocyte-specific disruption of soluble epoxide hydrolase attenuates abdominal aortic aneurysm formation: novel role of the liver in aneurysm pathogenesis"

**Figure S1.** Comparison of sEH expression in tissues and cells

#### BioGPS (GeneAtlas U133A, gcrma)

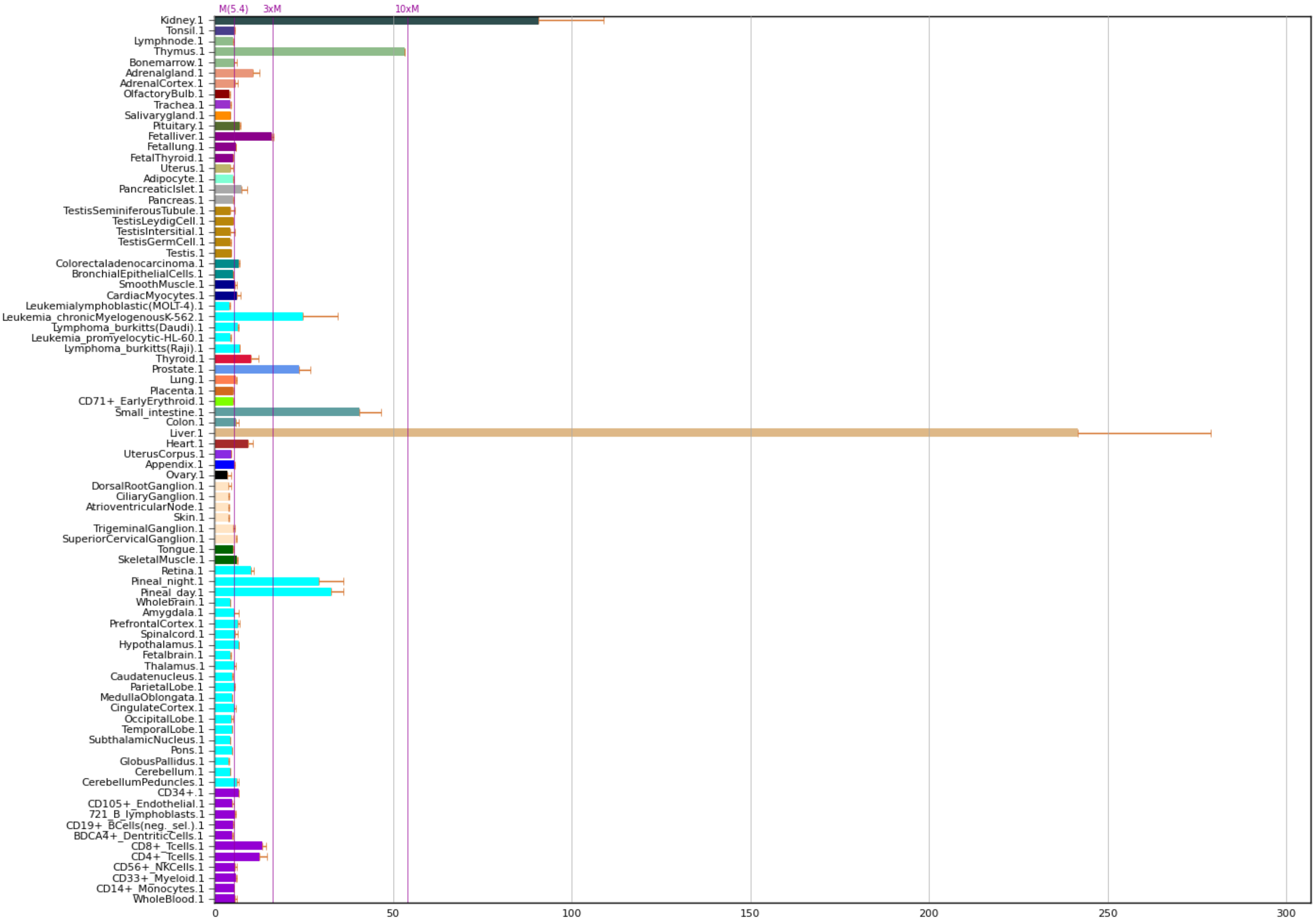

**Figure S2.** Cigarette smoke extract (CSE) increased expression of sEH, SAA and C3 in freshly-isolated liver slices *ex vivo*

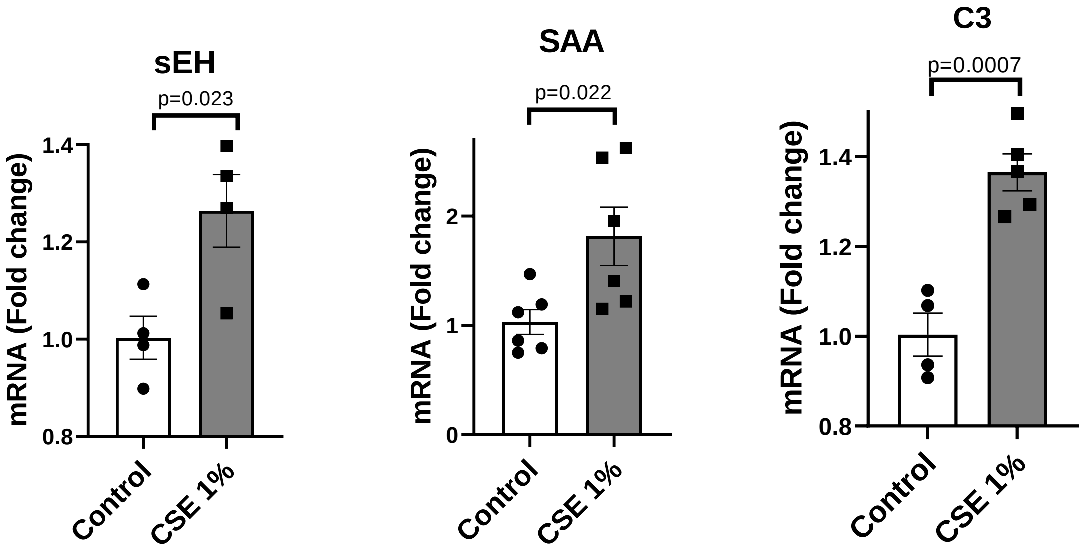

**Figure S3.** sEH inhibitor did not affect systolic blood pressure in AngII-infused mice

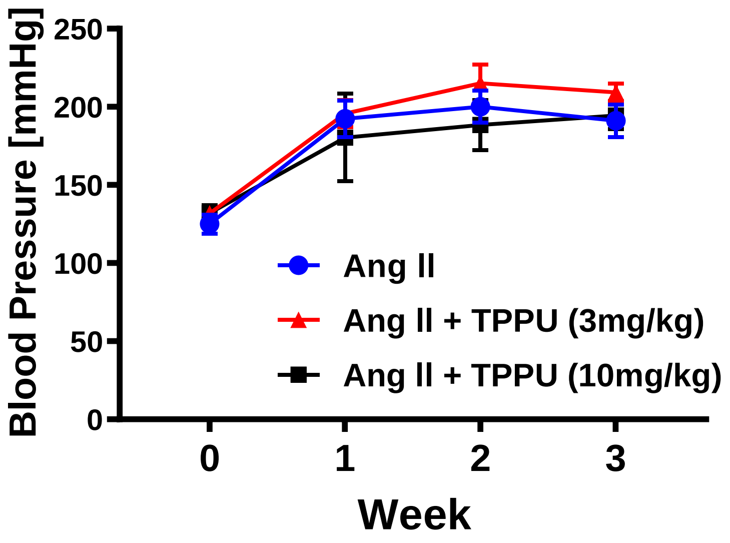

**Figure S4.** SAA mRNA expression in liver versus aorta: effects of AngII infusion and TPPU

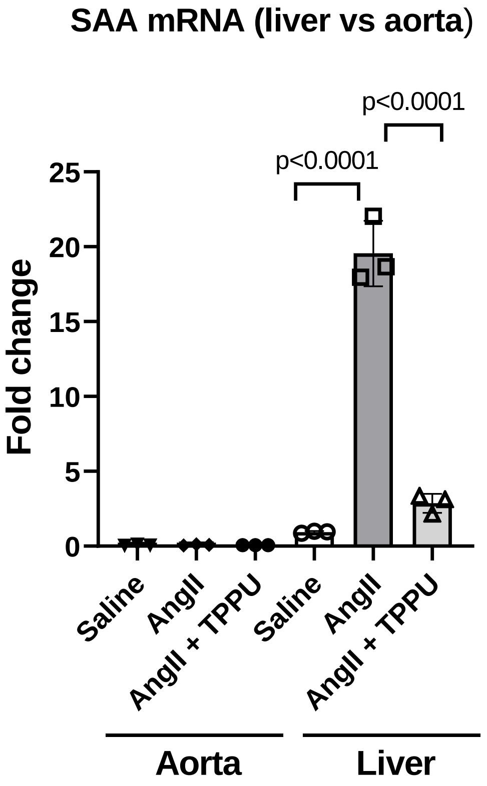

**Figure S5.** Fibrinogen mRNA expression in liver: effects of AngII infusion and TPPU

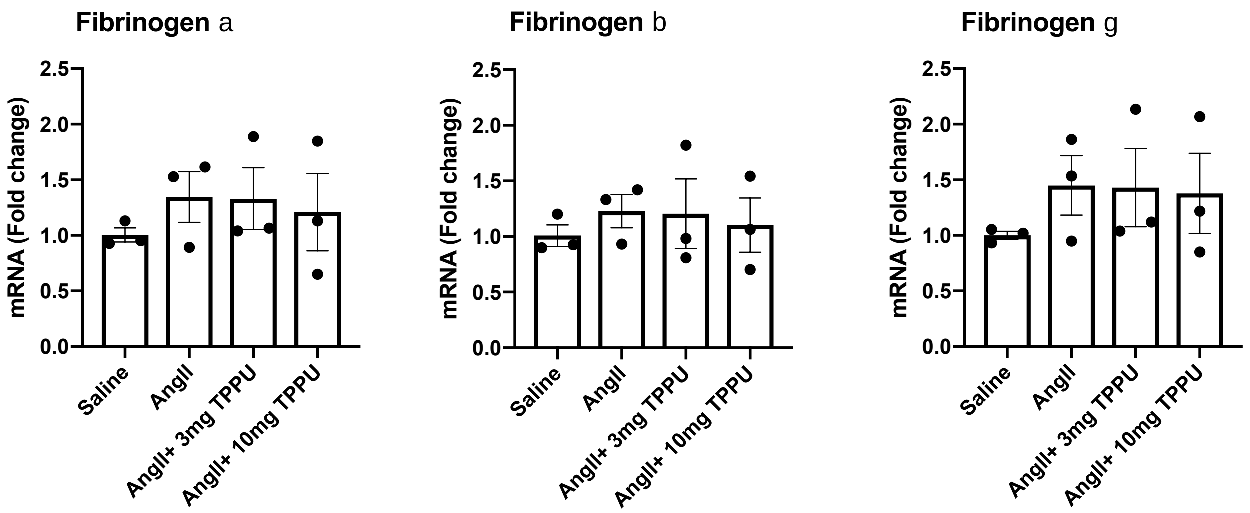

**Figure S6.** Validation of liver-specific sEH KO mice

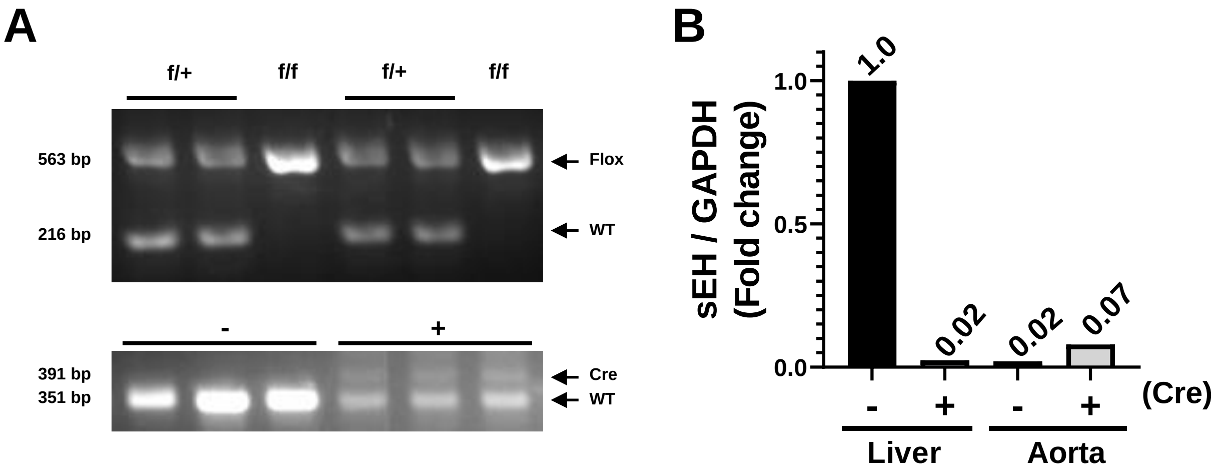

**Figure S7.** Liver-specific sEH gene deletion did not affect systolic blood pressure alteration in AngII-infused mice

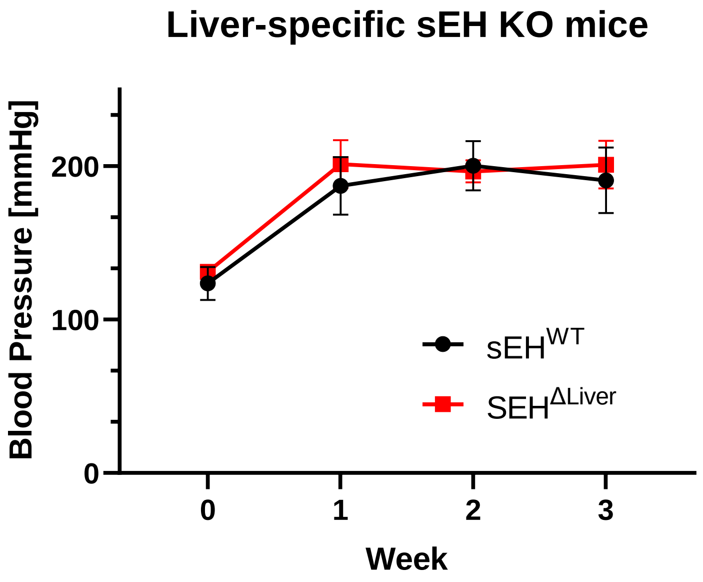

**Figure S8.** SAA expression in liver, plasma and aorta in $CaCl_2$-induced AAA model: effects of TPPU

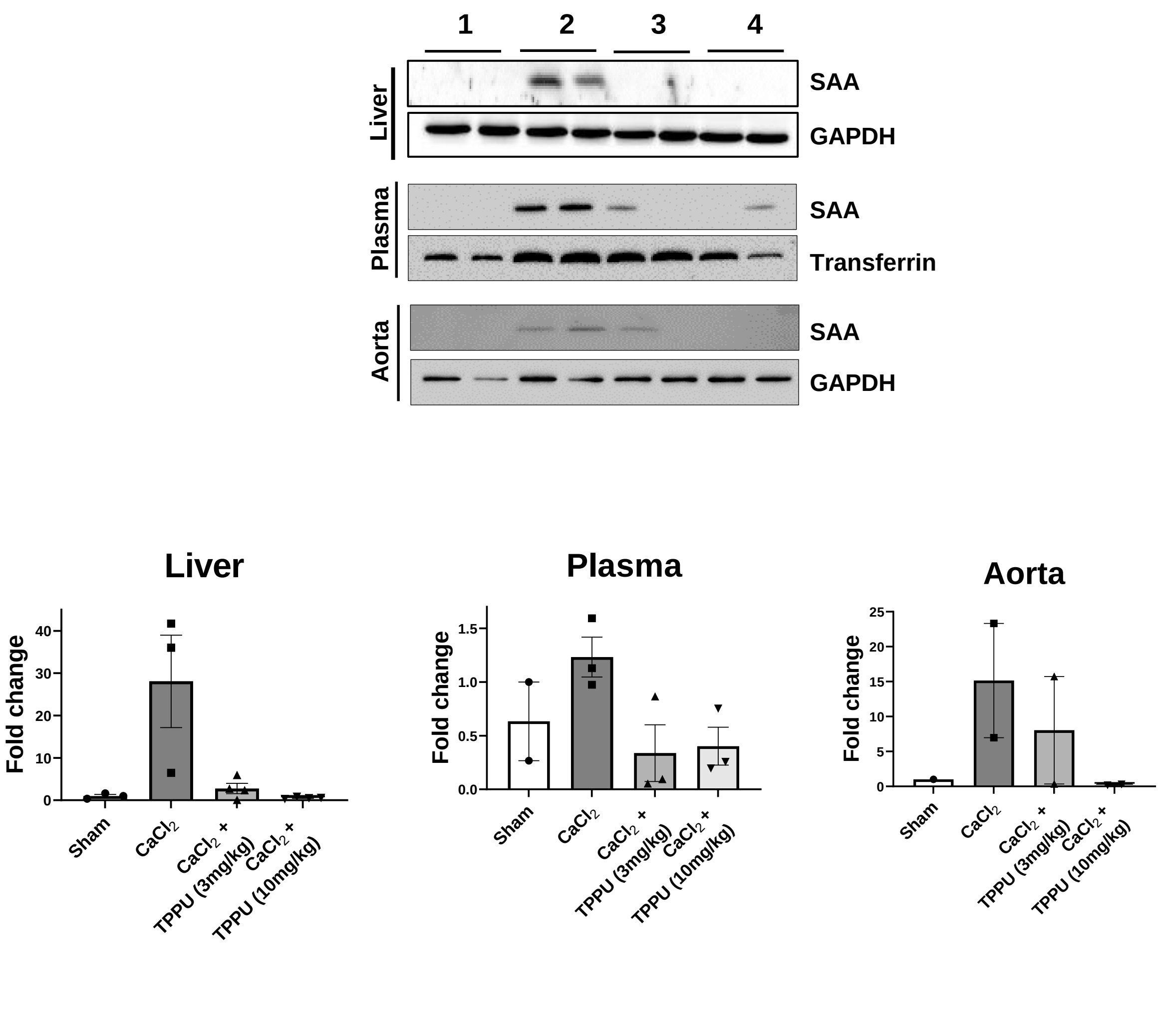

### Figure S9. Enriched GO categories: Upregulated

| GO ID | GO Name | C | O | Raw P-value | Adjusted P-value |
|---|---|---|---|---|---|
| GO:1900026 | positive regulation of substrate adhesion-dependent cell spreading | 34 | 4 | 1.9086e-8 | 0.0003 |
| GO:0045055 | regulated exocytosis | 747 | 8 | 5.6337e-8 | 0.0004 |
| GO:1900024 | regulation of substrate adhesion-dependent cell spreading | 47 | 4 | 7.2925e-8 | 0.0004 |
| GO:2001236 | regulation of extrinsic apoptotic signaling pathway | 146 | 5 | 1.2756e-7 | 0.0004 |
| GO:1902041 | regulation of extrinsic apoptotic signaling pathway via death domain receptors | 55 | 4 | 1.3888e-7 | 0.0004 |
| GO:0006887 | exocytosis | 849 | 8 | 1.5239e-7 | 0.0004 |
| GO:0016192 | vesicle-mediated transport | 1828 | 10 | 2.3545e-7 | 0.0005 |
| GO:0034116 | positive regulation of heterotypic cell-cell adhesion | 15 | 3 | 2.6892e-7 | 0.0005 |
| GO:2001233 | regulation of apoptotic signaling pathway | 372 | 6 | 4.492e-7 | 0.0008 |
| GO:0008625 | extrinsic apoptotic signaling pathway via death domain receptors | 79 | 4 | 6.0442e-7 | 0.001 |

### Figure S10. Enriched GO categories: <u>Downregulated</u>

| GO ID | GO Name | C | O | Raw P-value | Adjusted P-value |
|---|---|---|---|---|---|
| GO:0006735 | NADH regeneration | 25 | 6 | 2.609e-14 | 1.0774e-10 |
| GO:0061621 | canonical glycolysis | 25 | 6 | 2.609e-14 | 1.0774e-10 |
| GO:0061718 | glucose catabolic process to pyruvate | 25 | 6 | 2.609e-14 | 1.0774e-10 |
| GO:0061615 | glycolytic process through fructose-6-phosphate | 26 | 6 | 3.3973e-14 | 1.0774e-10 |
| GO:0061620 | glycolytic process through glucose-6-phosphate | 26 | 6 | 3.3973e-14 | 1.0774e-10 |
| GO:0019674 | NAD metabolic process | 66 | 7 | 5.5733e-14 | 1.4729e-10 |
| GO:0019362 | pyridine nucleotide metabolic process | 139 | 8 | 8.3267e-14 | 1.6505e-10 |
| GO:0046496 | nicotinamide nucleotide metabolic process | 139 | 8 | 8.3267e-14 | 1.6505e-10 |
| GO:0006007 | glucose catabolic process | 31 | 6 | 1.0814e-13 | 1.6933e-10 |
| GO:0006096 | glycolytic process | 73 | 7 | 1.1613e-13 | 1.6933e-10 |
